# Training load and body composition in adults practicing cyclical exercises

**DOI:** 10.1101/520189

**Authors:** Raquel Suelen Brito da Silva, Marizângela Ferreira de Souza, Matheus da Silveira Costa, Gisele Augusta Maciel Franca, Glêbia Alexa Cardoso, Joane Raquel Estrela Batista, Carolina Farias Arruda Lopes, Francisca Karinny Lemos Barbosa, Alexandre Sérgio Silva

## Abstract

Although meta-analyzes point to a weight loss of no more than 3 kg to exercise, body fat of the athletes are below of the population. Then training load may be a determining factor in body composition. This study verified if dose of physical training adopted by exercise practitioners is determinant in body composition. Was a cross-sectional retrospective study carried out with 122 individuals (45.8 ± 13.0 years, 50 men) who practiced cyclic exercises (running, walking or cycling) randomly recruited in six regions which the city was geographically divided. Caloric expenditure was estimated in the trainings based on the frequency, intensity and duration of the exercises and the body composition was assessed by electrical bioimpedance. The subjects practiced 4.3 ± 1.5 weekly sessions, with mean duration of 56.7 ± 28.2 minutes/session and caloric expenditure/day of 410.2 ± 384.1 kcal/day. Linear regression test revealed a negative correlation (p=0.000) between the mean daily expenditure and all measures of adiposity tested (absolute and relative body fat and visceral fat), and evidenced that the training load explains 56% of the proposed model. When adjusted for sex, the correlation remained in men and disappeared in women. Men’s with energy expenditure higher than 785 kcal/day presented lower fat stores than congeners with minor diary training load. Conclude that training load adopted by physical exercise practitioners is an influencing factor in the body composition of men, but not of women. Load adopted in conventional programs training seems insufficient to produce adequate body composition.

## Introduction

The energy balance between ingestion and caloric expenditure is an important modulator of body composition [1]. Currently, physical exercise is considered the most effective way to increase energy expenditure, which depends on the duration, intensity and weekly frequency of the sessions [2]. However, meta-analytic studies [3,4] and original data [5,6] have shown very modest results of physical training on weight loss of the no more than 3 kg of fat mass after program training of the 3 and 5x / week, with sessions of 45 to 60 minutes.

Among some attempts to explain the resistance to exercise-induced weight loss, the literature reports possible mechanisms that act as a compensatory adaptation of the organism that includes reduction of basal metabolism in people who perform exercises [7], increased nutrient intake [8], oxidative stress [9], hyperinsulinemia [10] and genetic profile [11].

Despite these compensation mechanisms, athletes present body fat percentage below of the normal range (11 to 16% for men and 14 to 18% for women) [12, 13]. Considering that athletes can spend between 2226 a 3877 kcal/day [14-16], it is plausible to assume that a high training load outweighs the compensation mechanisms that limit weight loss in community training programs, in which people exercise only 3 to 5 times a week and spend between 400 and 600 Kcal per session [5, 17].

Although it seems obvious, this relationship between training load and body composition is not determined. Previous intervention studies did not adopt different training loads to verify a possible greater efficacy of the programs with greater dose of exercise. Similarly, as far as we know, there are no studies that have attempted to answer this question retrospectively, comparing the energy demand of training loads of athletes and non-athletes who already practice physical training with their body composition, verifying if people who take higher loads have lower body fat stock.

In order to test this hypothesis, we conduced a retrospective population-based study with a representative sample of practicing cyclic exercises (running, walking and/or cycling) of a metropolitan city, which aims at determining the body composition of the subjects and verifying whether it shows any correlation with the training load adopted in their usual exercise routine.

## Material and methods

### Participants

The sample was selected by conglomerates, considering each of the six districts in which the city is divided geographically. From each district, two public places used to practicing physical exercise was randomly selected. The participants should be practicing of running, walking and/or cycling for at least one year, have a stable training routine over the weeks (frequency, duration and intensity), have not suffered interruption of the training routine for more than two weeks in the last three months and not have followed any protocol (nutritional or pharmacological, for example) intended for weight loss within the past 12 months. In case they performed another exercise modality, this could not exceed 20% of the weekly training load. Finally, participants could not exceed 200 minutes of moderate physical activity per week, in addition to the usual physical training.

### Recruitment of participants

Visits were performed during the hours of greater flow of practitioners (5:00 AM to 7:00 AM and 4:00 PM to 7:00 PM) between March 2016 and October 2018. Individuals were approached 1 in 3 at the time they were practicing physical exercise. Those who met the inclusion criteria were scheduled for interview and data collection.

### Dietary Profile

The food intake assessment was performed by applying the 24-hour Reminder [18], performed in duplicate, two days referring to food consumption on weekdays. The values of the nutrients of the diet were measured and evaluated by a nutritionist, using software Avanutri version 4.0 (AVANUTRI-RJ, Brazil).

### Body Composition

Body composition evaluation was performed using the Inbody bioimpedance (model 720, Biospace, Korea). Body weight, fat percentage, fat mass, visceral fat, waist-hip ratio and skeletal muscle mass were measured. For this evaluation, the volunteers followed some pretest procedures: being fasting, not having exercised moderate to high intensity in the 12 hours before the evaluation, not having consumed alcohol 48 hours before the test, not having ingested coffee, do not wear metallic jewelry or dental implants with metal during the evaluation and make use of light clothing during the procedure. Stature was verified by means of a Sanny^®^ professional stadiometer as recommended by the World Health Organization/WHO [19]. The body composition and nutritional data were collected at same week.

### Load Training

The training load was measured based on frequency, duration and speed adopted in exercises. According information of duration and distance covered by the participants, the speed of the training sessions was calculated. Those who did not know the exact distance traveled had a session monitored by means of a GPS. Knowing the modality, distance, speed and duration of sessions, was adopted the Compendium of Physical Activities proposed by Ainsworth et al. [20] to estimate the energy expenditure. Data were obtained in kcal by physical activity adjusted for body weight. The sum of the total caloric expenditure of all week sessions was divided by seven to adjust caloric expenditure to kcal/day.

### Statistical Analyses

After testing for normality and homogeneity, Pearson’s tests and linear regression were applied to verify the relashionship of the caloric expenditure/day on the body composition, using SPSS Statistics software (v. 24, *IBM SPSS*, Chicago, IL). In addition, the caloric expenditure in the exercise sessions was divided in terciles and ANOVA one way was used to compare the body composition of the participants in function of this categorization of caloric expenditure. Data were presented as mean and standard deviation of the mean or median and confidence interval (CI) 95%.

The study was approved by the Ethics Committee of the Health Sciences Center of the Federal University of Paraíba (CEP/CCS/UFPB), under protocol no. 0320/15.

## Results

The 122 participants had 45.8 ± 13.0 years. The men (n= 50) had body mass index /BMI of 26.1± 4.5 and women (n= 72) had 27.8 ± 5.7Kg/m^2^ They performed 4.3 ±1.5 sessions / week (56.7 ± 28.2 minutes/session). The mean daily caloric expenditure was 410.2 ± 384.1 kcal/day. Men obtained 641.2 ± 495.6 kcal/day (minimum and maximum values of 104.1 and 2142.1 kcal/day; women obtained 249.8 ± 135.2 kcal/day (47.0 to 653.3 kcal/day).

### Nutritional Profile

Caloric intake was 1927.2 ± 695.8kcal/day (28.4 ± 11.6 kcal/kg of body weight; 3.6 ± 1.6g/kg of carbohydrates, 1.4 ± 0.8 g/kg of proteins and 0.8 ± 0.4 g/kg of fats. When stratified by gender, the caloric intake was 2327.7 ± 735.4 kcal/day (32.9 ± 13.2 kcal/kg) for men and 1649.0 ± 510.3 kcal/day (25.2 ± 9.2 Kcal/kg) for women. Men and women presented 4.1 ± 1.9 and 3.3 ± 1.4g/kg of carbohydrates, 1.7 ± 0.9 and 1.2 ± 0.6g/kg of proteins, and 0.9 ± 0.5 and 0.7 ± 0.4 g/kg of fats, respectively.

### Food intake and body composition

Correlation tests ruled out the possibility that higher caloric intake, carbohydrate and dietary fats would account for the greater body fat composition. On the contrary, it was observed that higher caloric intake had a significant, however weak negative inverse correlation with fat percentage (p = 0.00, R = - 0.52) and absolute body fat weight (p = 0.00; R = -0.38). This same behavior happened when analyzes were performed for carbohydrate and fat intake, and in these cases, negative and weak correlations were also found for BMI and visceral fat. Was showed a positive and moderate correlation between caloric intake and daily caloric expenditure with exercise (p = 0.00, R = 0.53), with this same behavior being maintained for carbohydrate and fat intake.

### Training load, body composition and energy expenditure

The data presented in Table 1 confirm the hypothesis that the training load is an influencing factor on the body composition. There was a negative association between daily caloric expenditure in physical exercises and all measures of body composition related to adiposity. However, when the analysis was categorized by sex, it was observed that the correlation was maintained in men, and disappeared for all variables in women.

**Table 1.** Relationship between mean caloric expenditure per session and body composition of participants in the general population and stratified by sex.

|  | **General**<br>(47.0 - 2142.1<br>kcal/day) |  | **Men**<br>(104.1 – 2142.1<br>kcal/day) |  | **Women**<br>(47.0 – 653.3<br>kcal/day) |  |
| --- | --- | --- | --- | --- | --- | --- |
|  | **n= 122** |  | **n= 50** |  | **n= 72** |  |
| **Variables** | **Mean** | **R** | **Mean** | **R** | **Mean** | **R** |
| **Weight (kg)** | 71.6 ± 14.5 | 0.24 | 75.7 ± 14.5 | -0.25 | 68.8 ± 13.9 | 0.17 |
| **BMI (Kg/m<sup>2</sup>)** | 27.1 ± 5.3 | -0.21* | 26.1 ± 4.5 | -0.32* | 27.8 ± 5.7 | 0.08 |
| **Body fat (%)** | 31.7 ± 10.9 | -0.56* | 23.2 ± 9.0 | -0.49* | 37.6 ± 7.9 | -0.17 |
| **Body fat (kg)** | 23.3 ± 10.9 | -0.37* | 18.4 ± 9.7 | -0.41* | 26.6 ± 10.5 | 0.04 |
| **WHR (cm)** | 0.90 ± 0.1 | -0.32* | 0.88 ± 0.1 | -0.37* | 0.91 ± 0.1 | -0.05 |
| **Visceral fat (cm<sup>2</sup>)** | 90.4 ± 36.7 | -0.42* | 75.9 ± 41.9 | -0.43* | 100.6 ± 28.8 | 0.04 |
| **SMM (kg)** | 26.7 ± 6.1 | 0.45* | 32.2 ± 4.8 | 0.07 | 22.9 ± 3.2 | 0.38* |

Date: Between brackets below the general and stratified population are the minimum and maximum values of the caloric expenditure per day normalized by seven days week. Other data are mean and standard deviation of the mean. CE: caloric expenditure; BMI: body mass index; WHR: waist-hip ratio; SMM: skeletal muscle mass. \*: p<0.05; (Pearson Test)

Considering that the association was only found in men, a regression analysis was carried out only for this population, taking the fat percentage as dependent variable and the caloric expenditure/day as independent variable. This model confirmed that the caloric expenditure significantly influenced the fat percentage (p<0.00). The standardized beta coefficient indicated that caloric expenditure accounts for 56% of this variable among men.

Finally, the sample was divided into terciles (Table 2). It was verified that the subgroup of men in the upper tercile (>785 Kcal/day) presented the variables related to body fat significantly lower in relation to those in the lower tercile (caloric expenditure less than 398 kcal/day). Subjects in the middle tercile (between 398 and 785 kcal) were not significantly leaner than those in the first, nor in the third tercile. For women, the tercile distribution did not show differences between those with the lowest and the highest caloric expenditures per day.

**Table 2.** Body composition profile according to the caloric expenditure (in terciles) Date are mean ±SD. Mean values are caloric expenditure per session normalized by seven days week; CE: caloric expenditure; BMI: body mass index; WHR: waist-hip ratio; SMM: skeletal muscle mass. * = difference from the first tercile; # = difference from the second tercile (one way ANOVA).

|  | **General** |  |  | **Men** |  |  | **Women** |  |  |
| --- | --- | --- | --- | --- | --- | --- | --- | --- | --- |
| **CE/day**<br>**(kcal/d)** | **1st tercile**<br>**(< 247)**<br>**n=51** | **2nd tercile**<br>**(247-453)**<br>**n=36** | **3rd tercile**<br>**(> 453)**<br>**n=35** | **1st tercile**<br>**(< 398)**<br>**n=19** | **2nd tercile**<br>**(398-785)**<br>**n=19** | **3rd tercile**<br>**(> 785)**<br>**n=12** | **1st tercile**<br>**(< 188)**<br>**n=27** | **2nd tercile**<br>**(188-273)**<br>**n=20** | **3rd tercile**<br>**(> 273)**<br>**n=25** |
| **Weight (kg)** | 68.8±12.2 | 74.2±15.6 | 73.1±15.9 | 79.1±14.0 | 75.2±15.4 | 70.9±13.3 | 65.6±9.9 | 66.2±12.0 | 73.8±17.6 |
| **BMI**<br>**(Kg/m<sup>2</sup>)** | 27.3±4.4 | 28.2±6.0 | 25.7±5.5 | 27.5±4.7 | 25.9±4.3 | 24.2±4.4 | 27.1±4.4 | 26.9±4.8 | 29.3±7.3 |
| **Body fat**<br>**(%)** | 36.7±7.3 | 33.6±10.0 | 22.5±10.8*# | 28.0±7.8 | 22.2±7.6 | 17.3±9.5* | 39.2±6.5 | 35.4±8.8 | 37.7±8.3 |
| **Body fat**<br>**(kg)** | 25.6±8.0 | 25.6±11.5 | 17.4±12.0*# | 22.7±9.1 | 17.4±8.9 | 13.2±9.6* | 26.2±7.5 | 24.1±9.3 | 29.1±13.6 |
| **WHR (cm)** | 0.92±0.06 | 0.91±0.07 | 0.87±0.1*# | 0.92±0.1 | 0.89±0.1 | 0.84±0.1* | 0.91±0.1 | 0.91±0.1 | 0.91±0.1 |
| **Visceral fat**<br>**(cm<sup>2</sup>)** | 100.1±26.7 | 97.5±36.4 | 68.3±41.5*# | 93.9±36.9 | 73.4±39.5 | 51.4±42.5* | 100.3±22.2 | 93.4±32.4 | 106.9±31.9 |
| **SMM (kg)** | 23.6±4.6 | 26.8±5.8* | 31.2±5.4*# | 31.7±5.2 | 32.5±5.0 | 32.7±4.3 | 21.4±2.9 | 23.0±3.2 | 24.4±2.8* |

## Discussion

This study showed that men who spend higher calories in physical training have lower levels of body fat. The data obtained with women do not allow the same conclusion because this population has a small variability in caloric expenditure - while men’s energy expenditure varied from 104.1 to 2142.1 kcal/day, among women this range was only from 47.0 to 653.3kcal/day.

Compensatory adaptation to exercise has been pointed out as one of the explanations for limited weight loss found in prospective interventions studies [7] which some authors call of adaptive thermogenesis [21]. It is still suggested that oxidative stress, systemic inflammation explain this phenomenon [8]. Although physiological factors may explain exercise limitations to promote weight loss, the workout load is a factor that should not be disregarded, because empirical observation indicating that athletes present low fat stores. Although this relationship between training load and body composition seems obvious, at least as far as we know, this subject has been little explored.

The importance of considering this theme is that the limited ability of physical training to lose weight may simply be because the training load of 3 to 5 weekly sessions adopted in previous intervention studies may not be sufficient to provide clinically important weight loss. The studies of Li et al. [22] and Thomas et al. [23] correspond to the review studies that verified the relationship between training load and weight loss in intervention studies. However are based on studies with distinct populations and small sample sizes, which greatly limits these conclusions.

To investigate the participation of the training load, the studies can be designed methodologically in two ways, one of them with intervention studies with various training loads, to test a possible greater effectiveness of these loads in the loss of body fat. However, the difficulty in developing this type of study concerns the application of interventions in large populations involving various training loads for comparison purposes, which makes their viability quite difficult. Another possibility would be retrospective cross-sectional studies, verifying how people already submitted to various loads of training presents in terms of body composition. This second option is more practical and brings preliminary results that justify (or not) investment in intervention studies.

In addition to providing data from a systematic observation indicating that training load is a determinant factor in body composition, our data provide clues as to what would be the minimum daily training load to have adequate body fat stores. When the energy expenditure in the training was divided into terciles, our findings showed that only men’s that consumed more than 785 kcal / day presented significantly lower levels of body fat compared to those who had lower average daily training load. The demand is higher than that proposed by the American College of Sport Medicine [24], which indicates an energy expenditure of 300 kcal / day for reduction of total body mass and fat mass. Moreover, this is a training protocol far superior to the intervention studies that adopt 5 weekly sessions, with caloric expenditure of 400 –600 kcal/session [5, 17, 25], which adjusted for the seven days of the week, results in only 285.1 and 428.6 kcal/day, respectively.

Regression analysis showed statistically significant correlation between training load and fat stores, but also that the daily workout load represented 56% of the statistical model, which clearly indicates that, despite having the training load as an influencing factor among men, other factors are involved in the body composition response to physical training. In fact, this evidence is not surprising, since the literature has shown that several physiological [8], genetic [11] and behavioral [26] factors participate in the body composition.

Another important issue is that, among women, the training load was not a factor that influences in the body composition. One possible cause for such a finding is that women respond much less to training [22], as shown in several studies [5, 27, 28]. Another possible explanation is that the range of energy expenditure among women was very small, with maximum value of 653.3 kcal/day, while men burned 2142.1 kcal/day. An increase in the sample size, which is actually occurring in our study, can modify this scenario, as women who train with a higher caloric expenditure may appear in random sampling. Therefore, our data do not still allow us to determine whether the absence of correlation for women is due to physiological factors or to the training load.

Cross section studies have obvious limitations compared to prospective interventions. A single measurement of the variables was carried out, which makes us propose that future studies be done longitudinally to follow variations in these variables. Moreover, we consider only the energy expenditure in the training, being that the body has energy expenditure with other types of physical activities. Although we have adopted energy expenditure with other physical activities above 200 minutes of moderate physical activity per week as exclusion criterion, this daily energy demand, although small, can participate in the total energy balance, so we intend to consider it in future studies.

In terms of practical implication, we highlight the possibility that training programs aimed to weight loss in men should have the training loads reviewed. The main indication obtained from this study is that training programs with 3 to 5 sessions per week, with energy levels in the 200-300 kcal/week [29] may not be enough for those who need to obtain weight loss clinically important.

According to our data, men need to train seven times a week, spending 785 kcal in each session (equivalent to about 60 minutes with moderate to high intensity) to have adequate fat stores. Therefore, they are data that can modify the current paradigm regarding the relationship exercise and weight loss.

